# Metabolome Identification by Systematic Stable Isotope Labeling Experiments and False Discovery Analysis with a Target-Decoy Strategy

**DOI:** 10.1101/089904

**Authors:** Drew R. Jones, Xusheng Wang, Tim Shaw, Ji-Hoon Cho, Ping-Chung Chen, Kaushik Kumar Dey, Suiping Zhou, Yuxin Li, Nam Chul Kim, J. Paul Taylor, Udhghatri Kolli, Jiaxu Li, Junmin Peng

**Author notes:** These authors contributed equally to the work (Drew R. Jones, Xusheng Wang). Correspondence: Junmin Peng. **Abbreviations** MISSILE: metabolome identification by systematic stable isotope labeling experiments LCMS/MS: liquid chromatography-tandem mass spectrometry.

## Abstract

We introduce a formula-based strategy and algorithm (JUMPm) for global metabolite identification and false discovery analysis in untargeted mass spectrometry-based metabolomics. JUMPm determines the chemical formulas of metabolites from unlabeled and stable-isotope labeled metabolome data, and derives the most likely metabolite identity by searching structure databases. JUMPm also estimates the false discovery rate (FDR) with a target-decoy strategy based on the octet rule of chemistry. With systematic stable isotope labeling of yeast, we identified 2,085 chemical formulas (10% FDR), 892 of which were assigned with metabolite structures. We evaluated JUMPm with a library of synthetic standards, and found that 96% of the formulas were correctly identified. We extended the method to mammalian cells with direct isotope labeling and by heavy yeast spike-in. This strategy and algorithm provide a powerful a practical solution for global identification of metabolites with a critical measure of confidence.

## INTRODUCTION

Metabolomics aims to survey the global state of the small molecule profile in cells, tissues, and organisms. Metabolites are the substrates and products of myriad enzymatic reactions and are therefore considered to be direct readouts of biological activity. Many metabolites also function as building blocks, signaling factors, and molecular precursors which modify and regulate cellular components such as DNA, RNA, and protein. The human metabolome^1^ contains conventional cellular metabolites along with other chemicals derived from food, microbiota, and the environment. The role of the metabolome has been increasingly appreciated in both development and disease^2^. However, it is still a challenge to profile the complete metabolome due to the highly diverse chemical properties of small molecules and practical limitations of analytical strategies.

Liquid chromatography-tandem mass spectrometry (LC-MS/MS) is a prevalent method for global metabolome profiling^3^. Combining nanoscale LC with high-resolution MS leads to the detection of thousands of high-confidence metabolite features in a complex sample^4^. Numerous software programs have been developed for processing large-scale datasets^5-14^. Most of these programs share a common workflow, including feature detection, peak alignment, and relative quantification with semi-automated identification and/or laborious manual validation of selected peak features. Structural annotation of the selected features is typically achieved by searching against empirical MS/MS spectral libraries such as METLIN^15,16^, HMDB^1,17^, or NIST^18^.

Despite considerable progress in the development of software programs, identification of metabolites from untargeted studies remains a daunting task. One major limitation is that spectral libraries must be generated with synthetic standards. For instance, the NIST14 MS/MS database contains ~14,000 empirical MS/MS spectra, making it a precious but costly resource. To identify unknown metabolites, we need to consider potential compounds that may not be present in the spectral libraries. Theoretically there are more than 10^60^ compounds weighing 500 Da or less^19^; though the number of biologically relevant metabolites remains unknown. The largest public structure repository (PubChem) holds over 45 million entries^20^, though many of these compounds are synthetic or otherwise not applicable for biological studies^20^. Nevertheless, it would be difficult to expand the empirical MS/MS database approach to cover all metabolites across biological experiments. The other limitation is that none of the currently available programs estimate the false discovery rate for metabolite identification, a widely recognized limitation in the field^21^. With the accumulation of metabolite entries in spectral libraries, the probability of randomly matching experimental MS/MS spectra to the libraries is increasing. In addition, small molecules often yield much fewer product ions than large compounds (e.g. peptides in proteomics), exacerbating the problem of by-chance spectrum matches.

To address the limitations of spectral library searches and false discovery analysis, we propose a formula-based strategy for identifying metabolites, *metabolome identification by systematic stable isotope labeling experiments* (MISSILE), as well as a new program (JUMPm) for automated data analysis and false discovery evaluation. JUMPm is capable of processing unlabeled, partially labeled, and fully stable isotope labeled LC-MS/MS data. The MISSILE strategy substantially improves the confidence of formula assignment. We examined the MISSILE/JUMPm pipeline in yeast, extended it to mammalian cells, and validated it with a library of 500 synthetic compounds.

## RESULTS

### Theoretical evaluation of mass accuracy and isotope labeling on formula identification

We aim to unambiguously determine the chemical formula of a precursor ion and then search its MS/MS spectra against the metabolome database to identify candidate structures. To simulate this process, we searched the known masses of all unique formulas in the human metabolome database^1^ (HMBD, *n* = 8,255 up to 1,250 Da, **Figure 1a**) against a theoretical database of formulas (*n* = ~265,000,000, **Online Methods**). At a given mass tolerance, searches in the higher mass range showed a larger degree of ambiguous matches, consistent with the observation that molecular mass is exponentially correlated with the number of possible formulas^22^ (**Figure 1b, Supplementary Table 1**). We then simulated the effect of the MISSILE strategy which provides additional information on the stoichiometry of labeled atoms (C, H, N, O, P, or S, **Figure 1c**, **Supplementary Fig. 1**). Although each element alone provides limited discriminatory power, the combination of two (e.g. C and N) or more elements dramatically improves identification, resulting in a unique formula for almost all searches across the mass range. We therefore focused our efforts on achieving carbon and nitrogen labeling. This theoretical analysis demonstrates the potential advantage of the MISSILE strategy for formula identification.

**Figure 1.**
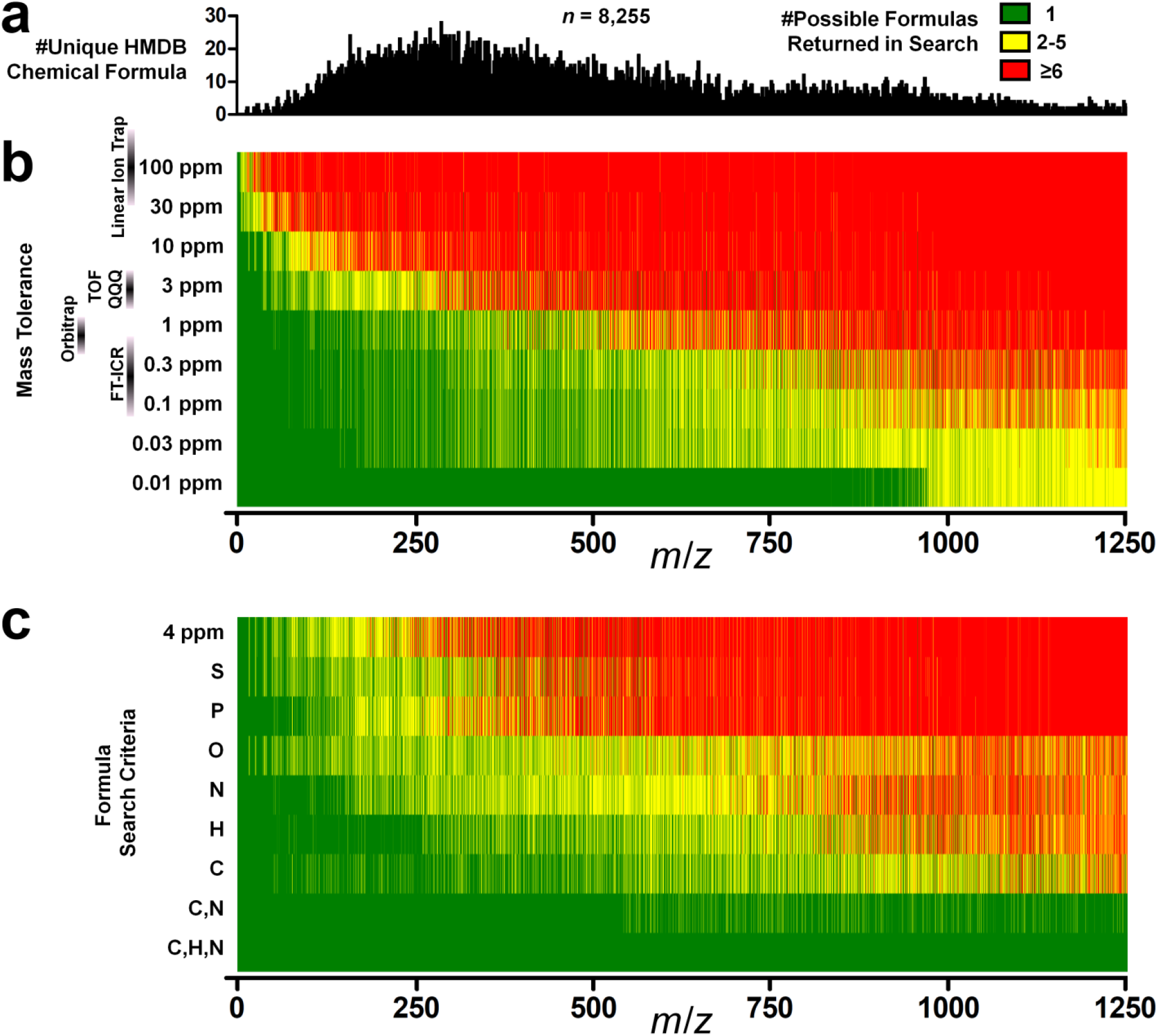
Simulated chemical formula searches with varying information. (**a**) Histogram of unique metabolite formula entries in the HMDB up to a mass of 1,250 Da. (**b**) Heat map of possible formula matches in the theoretical database as a function of mass tolerance (ppm) and precursor ion mass (Da. 8,255 HMDB mass inquiries). Colors indicate the number of possible formulas for any given search condition. (**c**) Metabolite formula searches restricted by known atom stoichiometry of assigned elements, with a mass tolerance of 4 ppm.

### JUMPm: automated metabolite formula determination and spectral matching

We developed JUMPm, a software program that automates the global analysis of unlabeled or stable-isotope labeled data using our formula-based strategy (**Figure 2a**). Our analysis uses metabolite chemical formulas to narrow down the possible structure candidates for a given peak and control the rate of false discovery. JUMPm accepts raw mass spectrometry data as input and then performs deisotoping, decharging, noise characterization, mass calibration, and feature detection prior to formula and structure searches (**Online Methods, and Supplementary Figs. 2-4**). For unlabeled or partially labeled samples, the program uses isotope pattern analysis to estimate the carbon atom number during the formula search (**Supplementary Figs. 5,6**). For labeled samples (i.e. MISSILE), JUMPm detects the labeled ion pairs with a Pairing score algorithm (Pscore, **Online Methods**), which considers three parameters including the unique isotopic mass defect of the ^13^C and ^15^N labels, the relative ion intensity, and the shape of the co-eluting peaks (**Figure 2b, Supplementary Figs. 7,8, Online Methods**).

**Figure 2.**
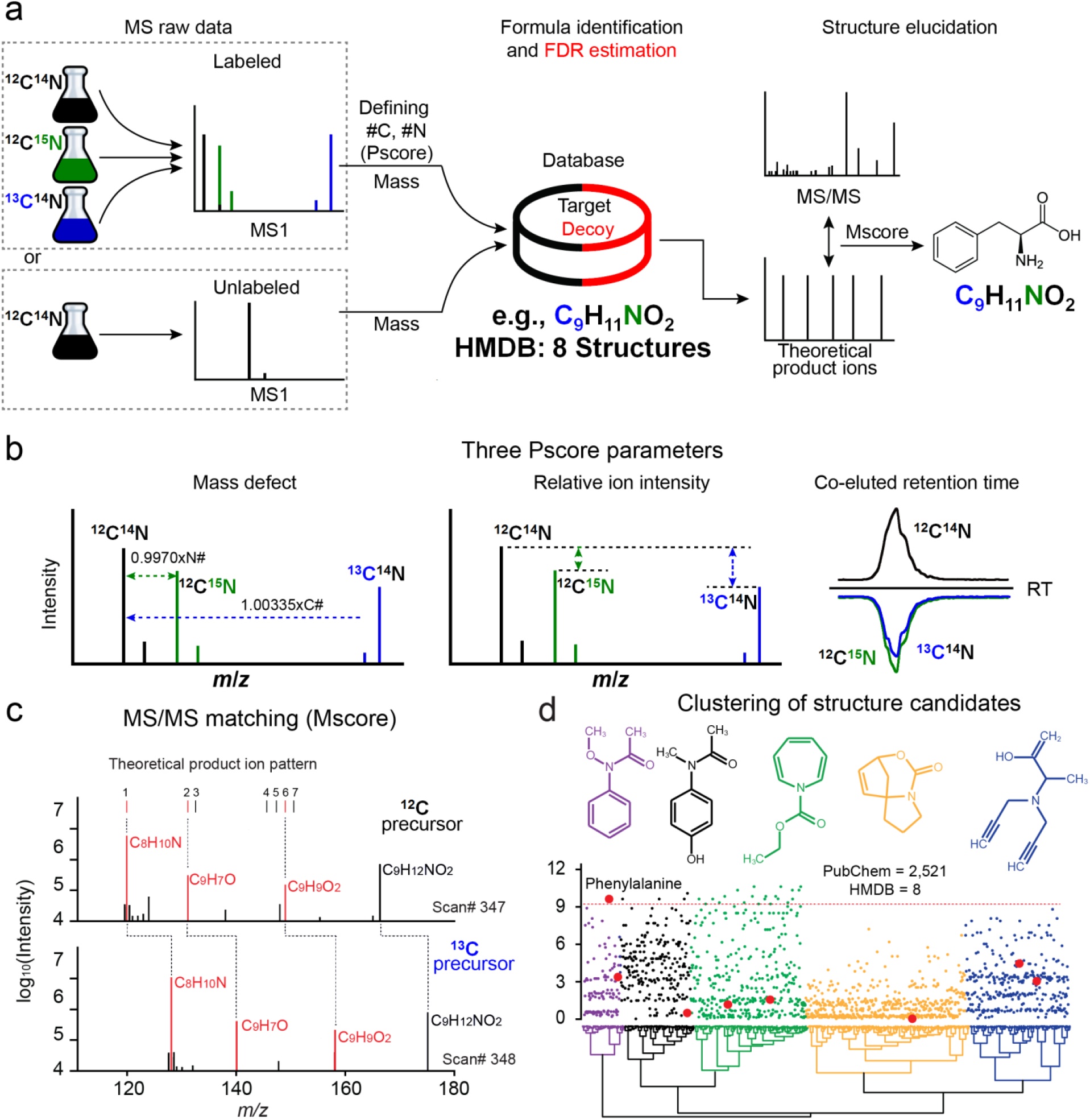
Overview of JUMPm workflow and an example analysis of phenylalanine in yeast. (**a**) Conceptual workflow for a stable-isotope labeling experiment with JUMPm data analysis. Yeast cultures were grown individually in various isotope labeled media conditions. Metabolites were extracted and combined in equal ratios to generate a mixed-label sample (phenylalanine MS1 shown). The three labeled peaks for a metabolite make up a “MISSILE” group. Full scan data are used for chemical formula determination and FDR estimation, while MS/MS data are used for structural identification of compounds with matching formulas. (**b**) The quality of each MISSILE is scored with three parameters. The Pscore is used to discriminate authentic MISSILEs from random matches. (**c**) For each MISSILE, the relevant MS/MS spectra are scored (Mscore) and annotated with the top match. MS/MS spectra from labeled metabolites include the extra mass of isotope labels in the product ions. (**d**) Hierarchical clustering of all structure candidates by predicted fragments for the example metabolite (HMDB candidates: large red dots; PubChem candidates: small dots). Representative structures from each colored group are shown. All candidates share the neutral formula C_9_H_11_NO_2_.

Once the metabolite formulas are identified, JUMPm finds any associated MS/MS spectra and searches them against a user-specified structure database (e.g. YMDB, HMDB, or PubChem), narrowing the search to only the candidates with that formula (**Supplementary Figs. 9,10**). This step significantly reduces the chance of a spurious annotation compared to traditional metabolite identification strategies such as accurate mass search, or spectral library search. JUMPm predicts the MS/MS fragments of database structures (with Metfrag^23^ or CFM-ID algorithms^24^) and ranks the candidates by a Matching score (Mscore) which compares the theoretical (*in silico*) and observed peaks (**Figure 2c, Supplementary Fig. 11, Online Methods**). A single chemical formula may have a large number of structural isomers (*mean* = 37 in PubChem) that may not be readily differentiated by MS/MS ions. For example, the formula C_9_H_11_NO_2_ yields 2,521 PubChem structural candidates that can be clustered into five analytical families based on shared fragments (**Figure 2d, Online Methods**). When searched against a curated database (e.g. HMDB), only 8 candidates are detected (large red dots in **Figure 2d**), with phenylalanine being the top hit. This analysis indicates that excessive search space increases the chance of spurious matches and reduces the possibility of identifying genuine metabolites, suggesting that the ideal database should be biologically relevant and contain expected compounds but be limited in size.

### False discovery evaluation with a metabolite target-decoy strategy

We implemented a target-decoy strategy to assess the degree of confidence in JUMPm metabolite identification. The target-decoy search strategy is a well-established method to analyze the false discovery rate (FDR) in other fields (e.g., proteomics^25,26^), so we developed a similar strategy to measure the rate of formula identification due to random chance. The target-decoy strategy typically uses a composite database containing half targets and half decoys such that the number of decoy hits (*n*_d_) are assumed to reflect the frequency of false matches. Therefore the FDR of target matches (*n*_t_) can be estimated by the equation (FDR = *n*_d_ / *n*_t_). The search results (target and decoy matches) are then filtered together by other parameters (e.g. mass accuracy and matching scores) to reduce the FDR to a user-defined level (**Supplementary Fig. 11c**).

The main challenge in applying this concept to metabolomics is to create decoys that adequately mimic targets yet are not valid hits, similar to reversed or randomized protein sequences in proteomics^25,26^. In chemical compounds, carbon, nitrogen, and oxygen follow the octet rule of chemistry, such that each atom has eight electrons in its valence shell (**Figure 3a, 3b**). There are rare exceptions to the rule^27,28^ (e.g., radicals or expanded octets), but we found that all of the HMDB entries follow the octet rule after accounting for these rare exceptions (**Supplementary Table 2**). To create decoy metabolites, we strategically violated the octet rule by adding one hydrogen atom to each formula in the database without changing the charge state of the entry (**Figure 3c**). These decoys mimic the mass distribution of targets, but can only be assigned due to by-chance matches. To test this strategy, we generated a negative control (null) dataset by shifting the ^12^C ion masses (+ 4.5 Da) of a raw file, creating essentially random masses. When searched against the composite target-decoy database, the target and decoy matches had an almost equal number (99%), indicating that all of the target hits from the null dataset are due to random matches (**Figure 3d, dashed line, Supplementary Fig. 12a**).

**Figure 3.**
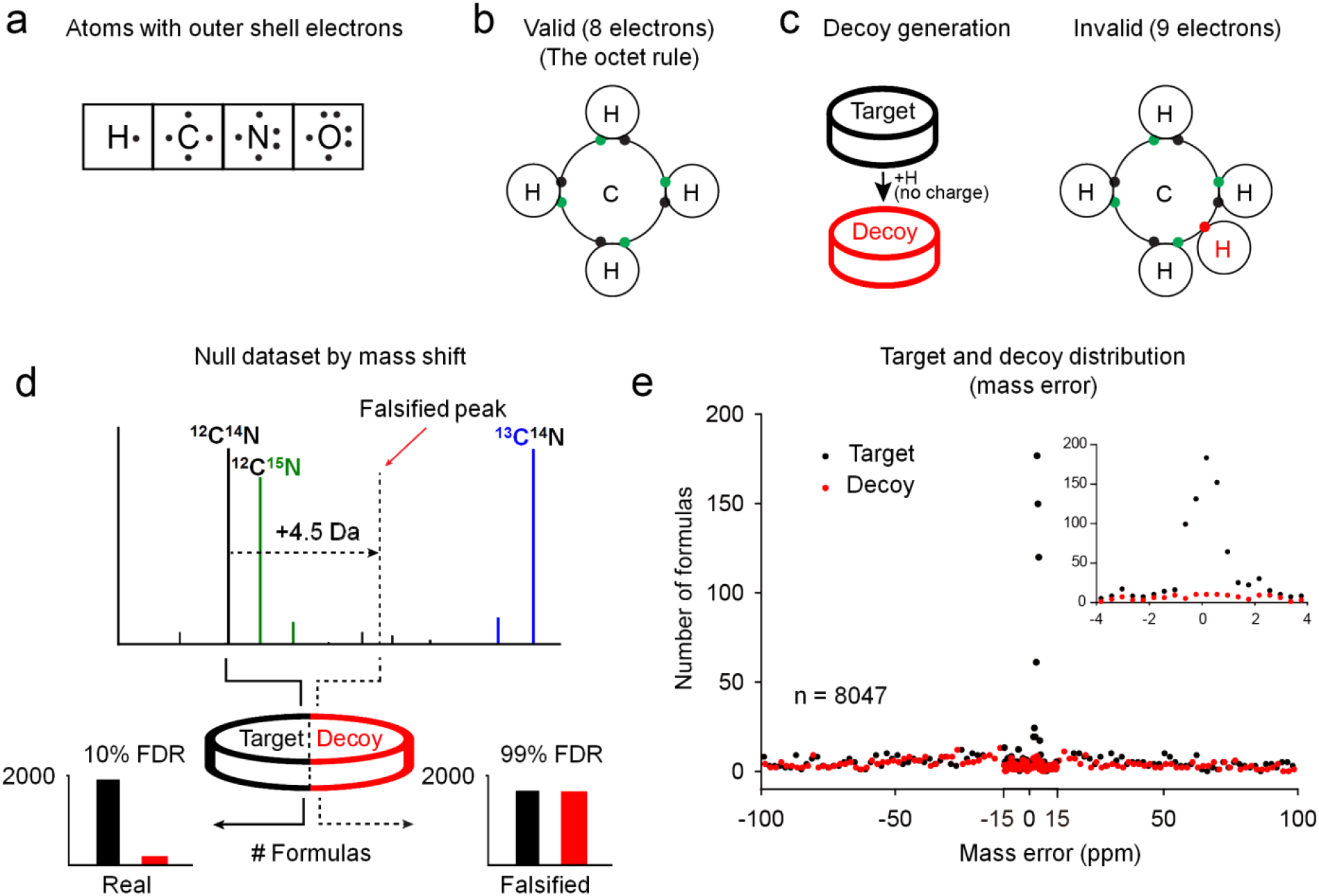
False discovery rate (FDR) estimation in JUMPm. (**a**) Common biological elements which follow the octet rule. Each element has a characteristic number of electrons available for bonding. (**b**) The valid Lewis structure for methane (CH_4_) shows shared electrons between hydrogen and carbon according to the octet rule. (**c**) Generation of decoy chemical formulas by computational addition of a hydrogen atom to each database formula, yielding an invalid structure without a change in the charge state. JUMPm treats all decoy formulas as neutral, ensuring that they are invalid. The impossible decoy structure for methane’s formula is shown. (**d**) FDR of authentic labeled yeast data (4%, n=102) and null data (~100%, n=6 targets, n=5 decoys). The relative ratio of decoy to target hits is an estimate of the FDR. (**e**) Histogram of target and decoy hits with respect to mass error during JUMPm search. Target and decoy hits are bins of 2 ppm across the mass error range (0.5 ppm within grey rectangle); a zoomed-in range is also shown.

In contrast, when we searched the authentic dataset (non-random input) (**Figure 3d, solid line**), there was a clear preference for the target database with an FDR of 10%, indicating that the pairing score algorithm accurately detected real isotope labels corresponding to real metabolite structures. For the authentic dataset, formulas with higher Pscores tended to have a lower FDR, suggesting a negative correlation between the Pscore and FDR (**Supplementary Fig. 12b**). When searching the LC-MS/MS data with a large mass tolerance (50 ppm), most of the targets were centered within a ± 2 ppm window, but the frequency of targets and decoys was equal outside of the window, indicating that those formulas were found due to by-chance matches because of the low mass accuracy (**Figure 3e**). We also inspected the formula distribution with respect to the mass defect of the labels (i.e., ^13^C and ^15^N, **Supplementary Fig. 12c**). Only target formulas were identified within ± 0.001 Da of the theoretical isotope mass difference (i.e. 1.00335 for carbon, 0.99703 for nitrogen). These results demonstrate that the target-decoy strategy is a powerful tool for assessing the confidence of identified formulas.

### Large-scale metabolome analysis in yeast by MISSILE/JUMPm

To explore the MISSILE/JUMPm pipeline for global metabolite identification, we carried out a comprehensive analysis of the yeast metabolome. First we characterized the labeling efficiency of the MISSILE strategy in yeast. The yeast strain grew at the same rate in the heavy isotope-labeled media (e.g. ^12^C^15^N or ^13^C^14^N) as in the standard unlabeled (^12^C^14^N) media (doubling time = 2.2 hr, **Supplementary Fig. 13a,b**). We then assessed the labeling efficiency by analyzing each yeast culture alone or mixed (^12^C^14^N + ^13^C^14^N; ^12^C^14^N + ^12^C^15^N, **Figure 4a-c**). There were no unlabeled peaks detected in the labeled samples, indicating complete labeling in yeast. Because the isotopic pattern of metabolites is largely determined by ^13^C, we observed that the ^13^C^14^N sample displayed a different (reversed) isotopic pattern from the unlabeled sample. Further analysis determined that the global labeling purity of the ^12^C^15^N and ^13^C^14^N isotope labels was at least 99% pure across the detected formulas (**Supplementary Fig. 14, Online Methods**).

**Figure 4.**
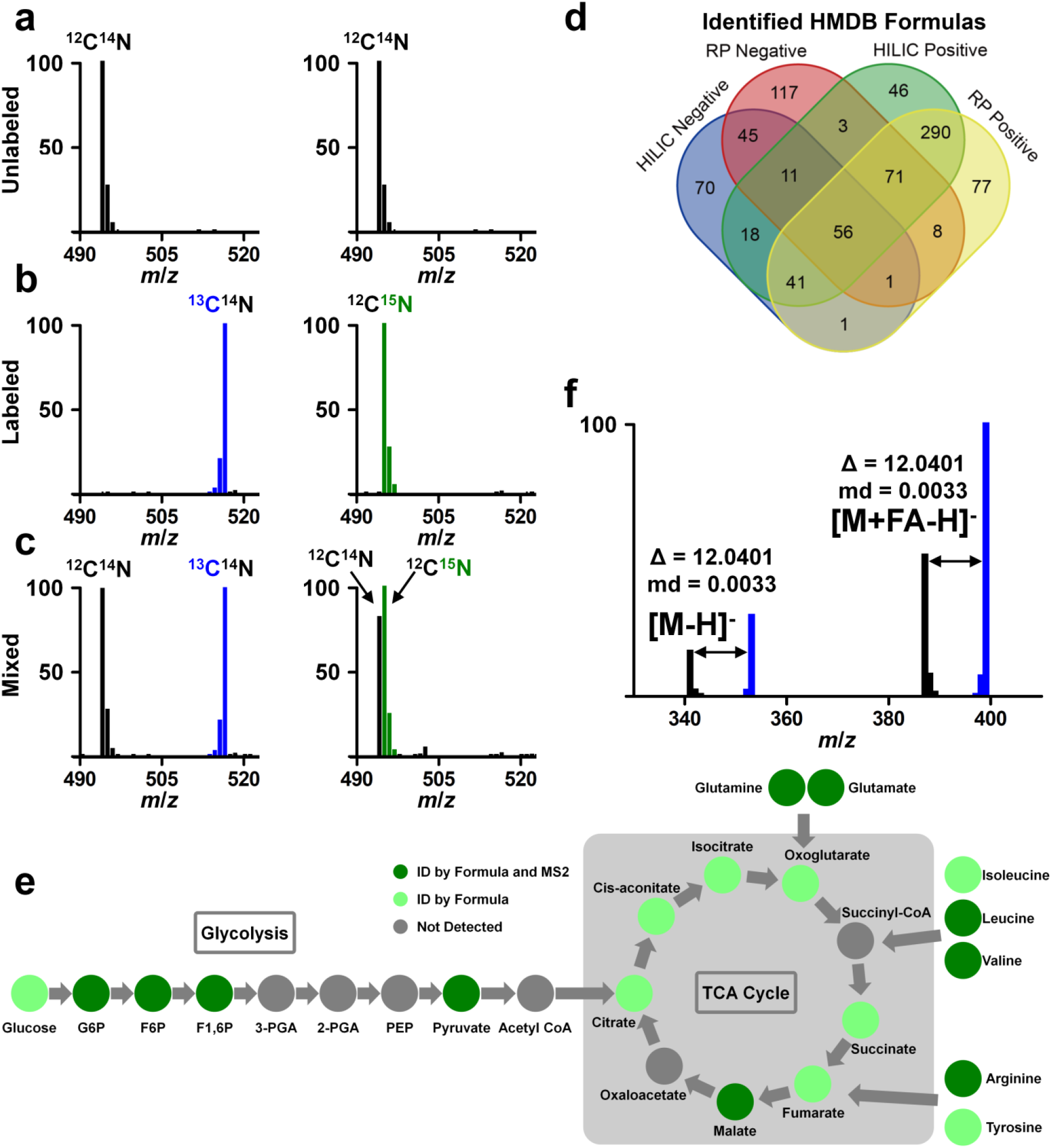
Large-scale metabolome analysis in yeast by MISSILE/JUMPm. (**a-c**) Labeling efficiency of stable isotopes in yeast with an example metabolite (494.3256 *m*/*z*). Labeled yeast samples were analyzed alone or mixed to assess the purity. (**d**) Overlap among triplicate analyses of labeled yeast metabolite extracts using C18 and HILIC columns in positive and negative mode, n=12 raw files searched by JUMPm (**Online Methods**). (**e**) Annotated map of the glycolytic and TCA metabolic pathways using JUMPm search results from yeast. Some compounds were identified by formula and the top MS/MS structure hit (dark green) or by formula only (light green). (**f**) MS1 spectrum of trehalose from a mixture of ^12^C and ^13^C yeast lysate. Compound identity was confirmed by external standards. [M-H]^-^ denotes the negative mode molecular ion, while [M+FA-H]^-^ denotes the negative mode formic acid adduct of trehalose. The formic acid adduct increases the apparent *m*/*z*, but does not affect the mass shift of the isotope label. Both isotope labeled pairs show a shift of 12 carbons.

Metabolites exhibit a diverse array of chemical properties, so we analyzed a mixture of all three yeast labels (^12^C^14^N + ^13^C^14^N + ^12^C^15^N) by four different LC-MS/MS conditions, including reverse-phase and HILIC chromatography in both positive and negative ionization modes, in triplicate (**Online Methods**). The four conditions were largely complementary with some overlap in identified metabolites (**Figure 4d, Supplementary Fig. 15**), totaling 2,085 metabolite formulas (10% FDR, **Supplementary Table 3**). This global, untargeted analysis covered 76% of the metabolites in the glycolytic pathway and TCA cycle (**Figure 4e**). To annotate the structures of the identified formulas, we matched the MS/MS spectra for each formula with various structure candidates across two databases; the yeast metabolome database (YMDB) and HMDB (**Online Methods**). We identified 892 metabolite structures in this study (**Supplementary Table 4**), which was limited by the lack of database candidates for many of the 2,085 formulas. We further examined the JUMPm algorithm at the fragment level by manually verifying the fragment formulas for a well-known compound, phenylalanine (**Supplementary Fig. 15a-d**). This structure annotation was validated by searching against the NIST14 MS/MS standard library, returning a probability score of 98.7% (R.Match: 997/1000) for phenylalanine.

We also investigated the impact of adducts on formula determination by JUMPm. Adducts are inorganic charge carriers (e.g., Na^+^, Cl^-^) or small acids/bases (e.g., formic acid, ammonia) in the sample matrix or LC mobile phase, which weakly bond with the analyte in the gas-phase during ionization. Adducts alter the *m*/*z* of the analyte, but do not contribute to the mass shift of MISSILE labeled metabolites (**Figure 4f**), and typically do not affect the MS/MS fragmentation pattern (**Supplementary Fig. 16a,b**). We implemented a function in JUMPm to consider the mass shift of user-defined adducts and identified 8 formic acid adducts from one raw file (**Supplementary Table 5**).

### Validation of MISSILE/JUMPm-identified metabolites with a synthetic standard library

To determine how reliably JUMPm identifies metabolites in untargeted metabolomics, we analyzed the heavy stable isotope labeled yeast extracts mixed with a commercially available metabolite library (500 synthetic standards with 394 unique formulas). First, we examined the quality of the library by dividing it into 20 cocktails for LC-MS/MS runs. A total of 337 (67%) of the standards were detected (S/N >100) in the LC-MS/MS runs with retention times recorded (**Supplementary Table 6**, **Online Methods**). Then we spiked the library into the ^13^C^14^N and ^12^C^15^N yeast metabolite extracts to recapitulate a MISSILE analysis (**Figure 5a**). JUMPm detected the MISSILE pairs arising from unlabeled standards co-eluting with corresponding labeled yeast metabolites. For instance, fructose 1,6-bisphosphate was identified based on two peaks (the ^12^C^14^N peak from the library and the ^13^C^14^N peak from the labeled yeast, **Figure 5b**) by JUMPm, matching the correct formula (C_6_H_14_O_12_P_2_). The annotation was further confirmed by the retention time of the standard in a separate run (**Figure 5c**). In another case, JUMPm identified and differentiated two distinct metabolites with the same formula but at different retention times; adenosine monophosphate (AMP, C_10_H_14_N_5_O_7_P) and dGMP (**Figure 5b-d**). Overall, we detected 91 standards with corresponding labeled yeast metabolites (4% FDR) in the spike-in experiment (**Figure 5e, Supplementary Table 7**). For 87 (96%) of these hits, JUMPm reported the same formula as the standard compound, in agreement with the estimated FDR. Further, the exact structural isomer for each formula was correctly annotated by JUMPm for 54% of the detected metabolites with data-dependent MS/MS scans. For those hits with different structures from the known standard, we found that the JUMPm annotation was typically a nearly indistinguishable isomer (e.g., xanthurenic acid vs. zeanic acid), which are generally not differentiated in a global LC-MS/MS analysis.

**Figure 5.**
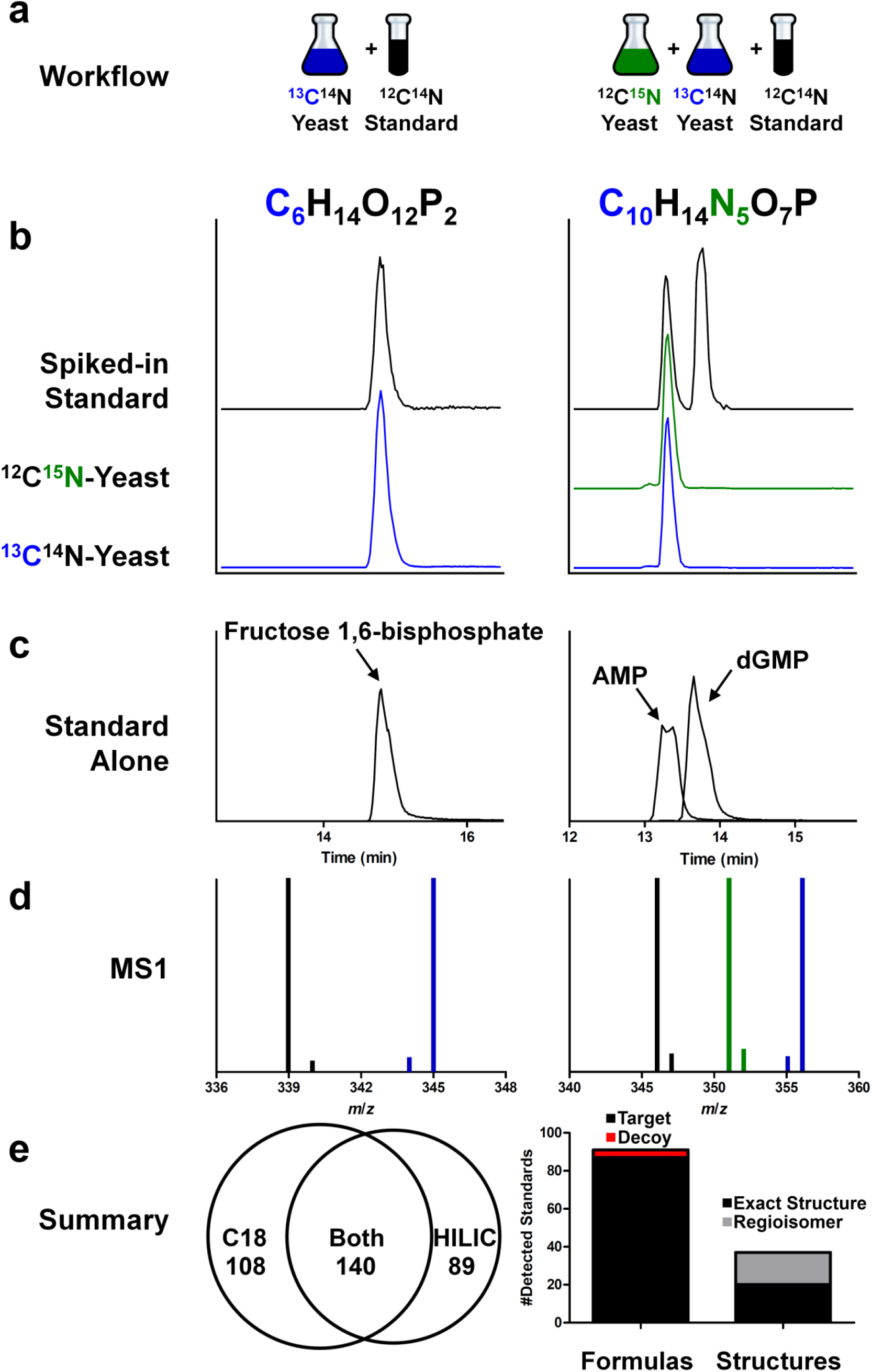
Validation of JUMPm-identified metabolites with synthetic standards. (**a**) Workflow for the spike-in experiment. Synthetic standards (black, n=500) were spiked-in with ^12^C^15^N (green), ^13^C^14^N (blue) labeled yeast extract, or both. (**b**) Extracted ion chromatograms of the standards and corresponding labeled yeast metabolites from the spike-in sample, showing the same retention time and peak shape. (**c**) Extracted ion chromatograms of the unlabeled standard in separate runs of the standard alone to confirm the retention time. (**d**) MS1 scan from the spiked- in sample to show the matching of unlabeled and labeled peaks. (**e**) Overall statistics on the detected standards by JUMPm. Left, the number of standards detected across the 20 cocktails by C18 and HILIC methods (**Online methods**). Right, the number of standards with corresponding labeled yeast metabolites for which JUMPm assigned the correct formula and structure for the spike-in analysis. Targets are correct formulas, while decoys are false formulas. Exact structures are from the known standards correctly identified by JUMPm, while regioisomers are JUMPm reported structures that are highly related to the known structure but differ slightly (e.g., glucose 1-phosphate vs. glucose 6-phosphate).

We also used the standards to evaluate the reliability of the widely used spectral library search strategy (e.g. NIST14 MS/MS) for metabolite identification. The NIST14 database contains spectra from 8,351 small molecules across 193,119 scans^18^. When searching the MS/MS spectra from the detected standards analyzed alone (n=337), NIST14 found the true formula for 50% of the standards (**Supplementary Fig. 17a-c**). Then we tried searching our structures (n=892) from the global yeast dataset (**Table 4**) with NIST14. Since the NIST14 MS/MS library is built from experimental spectral from unlabeled (^12^C^14^N) standards, MS/MS scans from labeled yeast parent ions (^12^C^15^N and ^13^C^14^N) served as negative controls (**Supplementary Fig. 17d**). About 22% of the NIST14 searches from our unlabeled yeast spectra gave the same formula as determined by JUMPm (**Supplementary Fig. 17e**). When we tried searching spectra from labeled parents against NIST14 (**Table 4**), none of the reported formulas matched the JUMPm formula. When JUMPm and NIST14 agreed on the formula for a given spectrum, these spectra had a statistically significant (p=0.0007) but small increase in their average score, similar to the difference observed between the true and false spectra of the standard compounds (**Supplementary Fig. 17c**). Therefore, spectral libraries (e.g., NIST14 MS/MS) can identify true hits, but are prone to high rates of false discovery (**Supplementary Fig. 17f**).

### MISSILE in other experimental systems

We further attempted the MISSILE strategy in human embryonic kidney 293 cells (HEK293), opening the way for labeled analyses in more complex samples. We identified HEK293 cell metabolites by directly labeling HEK293 cells with ^13^C-6-glucose in place of standard glucose or by spiking-in heavy labeled yeast extracts (**Supplementary Fig. 5b,c**). JUMPm is able to accommodate a variety of isotope labeling conditions, according to the user’s experimental design. We first mixed unlabeled and ^13^C-6-glucose labeled HEK293 cell extracts for LC-MS/MS analysis. Because there are multiple carbon sources in the cell culture media, we achieved ~50% purity among metabolites with glucose-derived ^13^C atoms. Therefore, we used the partial labeling search option in JUMPm, which enables JUMPm to detect partially labeled ion clusters and use isotope pattern simulation to determine the number of carbon atoms in the formula. Using this function, JUMPm identified 71 metabolite formulas from directly labeled HEK293 samples (**Supplementary Table 8**). We were also able to identify 219 unique formulas and 197 structures from unlabeled HEK293 cells by spiking-in heavy metabolites from yeast cells with an FDR of 3% (**Supplementary Table 9**).

## DISCUSSION

MISSILE/JUMPm is a comprehensive strategy for global identification of metabolite formulas and structures. Many isotope labeling conditions are possible, likely in any organism that can be grown with synthetically defined (SD) media, or where the carbon and/or nitrogen sources can be efficiently replaced with isotope labeled sources. Alternatively, a wide range of samples can be analyzed with the spike-in strategy, as long as the metabolites are found in both yeast and the system being studied. The use of isotope labeled samples increases the confidence of metabolite identifications in untargeted experiments and helps exclude false matches in MS/MS database searches by only considering candidates with the specified chemical formula. A chemical is also a useful annotation for unknown structures that can be referenced in subsequent studies. Tandem MS data (MS/MS) can be used to probe the substructure of novel metabolites, providing hypotheses for the chemical structure. Therefore the user’s choice of structure database will depend on the analyzed samples and experimental goals. We recommend searching HMDB for biological studies and routine identifications while PubChem may be useful for novel structure/similarity searches. Custom structure databases are also easily accommodated (**Online methods**). For example, results from MISSILE samples can be used to generate custom libraries for future analyses of unlabeled samples in the same experimental system.

Stable isotope labeling can improve the accuracy of metabolite formula identifications^29-34^ by greatly reducing the pool of candidate structures during annotation. We used isotope labeling methods to exploit the light-vs-heavy mass difference to experimentally determine the partial stoichiometry of a metabolite’s chemical formula. When combined with accurate mass we were able to determine a unique chemical formula. Global formula determination will expedite identification of known compounds, and aid in the discovery of unknown structures. Further, we automated the analysis of MISSILE data by developing the JUMPm software which can derive formulas and compare MS/MS spectra against the theoretical fragments of any database structure^23,35^.

Despite the advantages of the MISSILE method, there are several limitations of the approach. Inherently, a chemical formula is not a unique designation because many possible isomers may exist for a single formula. Exhaustive *de novo* structure generators routinely identify thousands to millions of potential structures for a given formula depending on the number of atoms^36^. Between constitutional and stereoisomers, the latter provide the biggest challenge for identification by tandem mass spectrometry. Constitutional isomers may have very different structures despite sharing the same chemical formula, and therefore typically give rise to unique MS/MS fragments. In contrast, stereoisomers typically generate the same MS/MS fragments, making it impossible to differentiate these candidates with LC-MS/MS alone. These limitations are also shared by traditional metabolite identification methods. These challenges may be addressed by reporting metabolite “groups”, similar to proteomics, and by improving the accuracy of fragment prediction algorithms. The exact structure of crucial hits may also be identified as needed by other analytical techniques including NMR.

Coverage is a critical issue for large-scale metabolome analyses. The number of identified metabolites will be affected by analytical coverage i.e., how many peaks are detected by LCMS, and by bioinformatic coverage i.e., and by how many authentic peaks are detected and annotated by JUMPm. In this study we used commercially available standards and spectral libraries based on human metabolites. When applying these tools to the analysis of yeast samples, we observed a significant drop in the identification rate. JUMPm is agnostic with respect to formula identification, so it is not affected by the selection bias found in empirical libraries. Analytical coverage is still a major challenge for metabolomics. While the four LCMS methods employed in this study were complementary, we still did not detect a significant number of the synthetic standards. After manual inspection we found that some of the missing compounds were present as dimers (multimers), in-source fragments, or other more complex forms. We also found that some previously detected standards were no longer observed when we spiked-in the highly complex yeast metabolite extracts, reducing the number of detected standards with corresponding labeled yeast structures. These results also point to a large number of yeast metabolites that are not currently annotated in any structure database (e.g., YMDB). In-depth analysis of multiple sample types will improve database coverage and help identify novel structures.

In summary, we have developed the MISSILE strategy along with JUMPm for the automated global analysis of metabolite formulas and structures in untargeted studies. JUMPm processes unlabeled, partially labeled, and fully labeled data, making it applicable to most systems of interest. We also introduced a novel target-decoy method for metabolomics, which estimates the FDR for identification, ensuring high-confidence results. We evaluated our strategy and software with a variety of datasets including a standard library, demonstrating that MISSILE/JUMPm is a simple and robust solution for untargeted metabolomics studies (freely available).

JUMPm download link:

<u>https://docs.google.com/uc?id=0B-8nCkZ-m2LhbHYwSm9vQ2RIMHM</u>

Database download link:

<u>https://docs.google.com/uc?id=0B-8nCkZ-m2LhbUczaDBjbDE4a2s</u>

## ACKNOWLEDGEMENTS

This work was partially supported by NIH grants AG047928, AG053987, GM114260 (J.P.), and ALSAC (American Lebanese Syrian Associated Charities). The MS analyses were performed in the St. Jude Children’s Research Hospital Proteomics Facility in the Hartwell Center, partially supported by NIH Cancer Center Support Grant (P30CA021765).

## AUTHOR CONTRIBUTIONS

J.P., D.R.J, and X.W. designed the research; D.R.J., P.C.C., K.K.D., N.C.K., J.P.T., U.K. and J.L. performed the labeling experiments and MS analysis; X.W., T.S, J.H.C., S.Z., and Y.L. developed the JUMPm computer software; X.W., D.R.J., and J.P. analyzed the data; D.R.J., X.W., and J.P. prepared the manuscript.

